# Modeling Protein Association from Homogeneous to Mixed Environments: A Reaction-Diffusion Dynamics Approach

**DOI:** 10.1101/2021.01.25.428073

**Authors:** Suraj Kumar Sahu, Mithun Biswas

## Abstract

Protein-protein association *in vivo* occur in a crowded and complex environment. Theoretical models based on hard-core repulsion predict stabilization of the product under crowded conditions. Soft interactions, on the contrary, can either stabilize or destabilize the product formation. Here we modeled protein association in presence of crowders of varying size, shape, interaction potential and used different mixing parameters for constituent crowders to study the influence on the association reaction. It was found that size a more dominant factor in crowder-induced stabilization than the shape. Furthermore, in a mixture of crowders having different sizes but identical interaction potential, the change of free energy is additive of the free energy changes produced by individual crowders. However, the free energy change is not additive if two crowders of same size interact via different interaction potentials. These findings provide a systematic understanding of crowding influences in heterogeneous medium.

## 1. Introduction

Protein-protein association is key to several biological processes like enzyme catalysis [1] and polymerization (for example, amyloid fibril formation) [2]. *In vivo* protein association reactions are impeded by the volume occupied by other proteins, enzymes, DNA-RNA, metabolites and osmolytes which can occupy approximately 30% of cellular volume. Recent evidences suggest that the cellular environment, apart from excluding available volume, plays a role in the association reaction by its physical [3–6] (like size, shape of molecules) and chemical [7–10] (like interaction potential) composition. In particular, weak soft interactions like hydrogen bonding and hydrophobic interactions are ubiquitous in cell and might play a more dominant role than exclusion interactions.

The first investigation of the role of cytoplasmic environment on the biophysical properties of proteins was carried out by Ogston[11] and Laurent[12]. Thereafter, Minton and others[13–33] have extensively studied how crowding conditions might affect protein-protein interactions in cellular media. In early *in vitro* experiments the cytoplasmic environment used to be mimicked by adding macromolecules such as dextrans, ficolls, and PEGs in varying concentrations. The observations many of these experiments showed that excluded volume interactions play the dominant role in crowding, consistent with the theoretical models [3, 34, 35]. In contrast, many experiments and simulations employing protein crowders showed opposite effect, that is, weak non-specific soft interactions play major role in crowding [8, 23–25, 36, 37]. ‘Soft’ interactions (like electrostatic, hydrophobic and van der Waals interactions) are long-ranged in comparison to ‘hard’ repulsive interactions which are dominant only at short distances. Quantifying the influence of soft interactions is challenging, since the forces can be both attractive and repulsive. Nevertheless, the influence of soft interactions has been evaluated by measuring the second virial coefficient [38–40], using ^15^N-relaxation experiment[41, 42] and in computational models [24, 43, 44].

Minton[13] first formulated a theoretical view of crowding influences on protein-protein interactions based on the scaled particle theory (SPT) [45–47]. Consider a binary association reaction between two protein species *A* and *B* producing *C*, that is, *A* + *B* ⇌ *C*. Let another species *D* is present in the reaction volume that influence the thermodynamics of binary association. Then, it is straightforward to show that the free energy change Δ*F* of the reaction in presence of crowders relative to the crowder free solution (that is when crowder packing fraction *ϕ*=0) is [6]

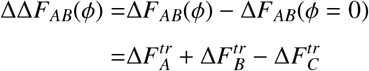

where 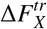 is the free energy to transfer species *X* (*X* ∈ (*A, B, C*)) to a crowded solution consisting of only species *D* from infinite dilution. In SPT, crowders are modeled as hard convex particles and the free energy to transfer any particle 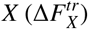 from a dilute solution to a crowded environment is calculated as [3]

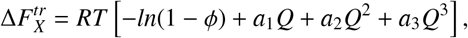

where *ϕ* denotes the volume fraction of the fluid spheres, 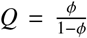 and *a*_1_*, a*_2_*, a*_3_ depend on the radii of the crowder *D* and particle *X*. In general, 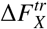 is estimated using Monte Carlo simulation or Widom’s particle insertion method and is used to obtain ΔΔ*F*_*AB*_(*ϕ*) [48]. But, here we adopt a direct approach to calculate ΔΔ*F*_*AB*_(*ϕ*) by following the reaction equilibrium under crowded and dilute conditions (detailed later).

A crowded milieu in the cytosol is a mixture of different kind of molecules. Evidences suggest that the influence of a mixed solution of crowders on protein stability and binding can be different from that observed in presence of its individual components [20, 49–51]. Thus, in order to understand protein association equilibria in a heterogeneous medium, it is essential to understand the role of mixing on crowding induced stabilization. In this work, we systematically study crowding effects on the thermodynamics of a binary association reaction using computer simulations and compare with the analytical result (SPT). Crowders are modeled with varying size, shape and pairwise interaction potential between constituent species. We show that crowding can both stabilize and destabilize the reaction depending upon the nature of interaction (attractive or repulsive). Further, we mix crowding species having different sizes and interaction potentials and discuss the role of mixing conditions on overall stability.

## 2. Methods

To investigate crowding effects in a cytosolic environment, consider the reaction *A* + *B* ⇌ *C* in equilibrium, that is, species *A, B* diffuse in a periodic box and come in contact with each other to form a product *C*. The reaction can occur with crowder molecules *D* in the box or at infinite dilution (no crowders). Using the equilibrium concentration of different species *n*_*i*_ (*i* = *A, B, C*) the free energy difference for the forward reaction at infinite dilution and with crowders can be directly evaluated as

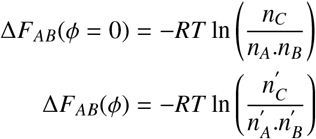

where 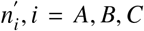 denote equilibrium concentration of species in presence of crowders.

In this work, the reaction was set up by employing a Brownian dynamics reaction-diffusion simulation framework as implemented in the *ReaDDy* package.[52] In *ReaDDy*, proteins and crowders are modeled as coarse-grained spherical particles having certain radii (see Table 1 for details). Although the severe coarse-graining results in loss of atomistic details crucial for several biological functions, it allows us to study the systematic trends with variation of crowding conditions, which would otherwise be highly computationally demanding. Non-spherical proteins or macromolecules were modeled as a group of particles having certain topology (shape).

**Table 1:** Parameters used in *ReaDDy* simulations.

| Parameter | Value |
| --- | --- |
| Box size (nm<sup>3</sup>), $b$ | (106.2)<sup>3</sup> |
| Temperature (K), $T$ | 300 |
| Number of A particles, $N_A$ | 500 |
| Number of B particles, $N_B$ | 500 |
| Water viscosity (kg/nm/ns), $\eta_0$ | 10<sup>-21</sup> |
| Radius of A (nm), $r_A$ | 3.0 |
| Radius of B (nm), $r_B$ | 3.0 |
| Radius of C (nm), $r_C$ | 3.78 |
| Radius of D (nm), $r_D$ | 5.0, 2.5 |
| Diffusion constant of A ($\mu\text{m}^2/\text{s}$), $D_A$ | 73.2 |
| Diffusion constant of B ($\mu\text{m}^2/\text{s}$), $D_B$ | 73.2 |
| Diffusion constant of C ($\mu\text{m}^2/\text{s}$), $D_C$ | 58.1 |
| Diffusion constant of D ($\mu\text{m}^2/\text{s}$), $D_D$ | 43.9, 87.9 |

For the cytoplasmic model of *M. genitalium*, the Stokes radius of proteins vary in the range 1.6-12.7 nm with the most abundant protein size being ~2.5 nm [29]. In this study, species *A* and *B* have a radius of 3 nm each. Although having the same size, different nomenclature is adopted since protein monomers of similar sizes may self- or hetero- associate to form a product complex. For example, many of the *α*, *β* and *γ* crystallin proteins found in mamallian lens have similar sizes and can undergo self- or hetero- association [53]. Product and reactant size and shape can also dramatically affect crowder induced stabilization [33, 54]. But, variation of these parameters is planned to be part of a forthcoming report, focusing only on the variation of crowder properties in this work.

Here the solvent was treated implicitly by using the diffusion constants of proteins and crowders in water as input parameters. The particles interact with each other by a pair-wise interaction potential which can be set either purely repulsive (for example, half-harmonic) or attractive and repulsive (for example, Lenard-Jones(LJ)). The half-harmonic potential *V*_*hh*_(*r*) is modeled as

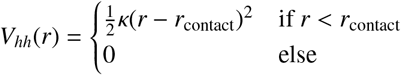

where *κ* is the force constant for the harmonic potential, *r* is the distance between *i*-th and *j*-th particle and *r*_contact_ is the corresponding contact distance, that is the distance between particle centers when the surfaces touch each other. The LJ potential has the form

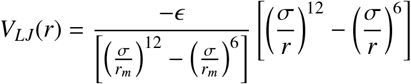

where *σ* is the distance where *V*_*LJ*_(*r*) = 0, *r*_*m*_ is the distance at which *V*_*LJ*_(*r*) reaches minimum and *ϵ* is the well-depth such that *V*_*LJ*_(*r*_*m*_) = −*ϵ*. The LJ potential is cutoff to zero at a distance of 2.5*σ* and is shifted to avoid a discontinuity. To compare the influence of repulsive and attractive potentials, we have set *σ* = *r*_contact_ (see Fig. 1). The well-depth *ϵ* of LJ potential was chosen to be 4 kJ/mol (1.6 *k*_*B*_*T*) ensuring simulation conditions explore a single phase region [44].

**Figure 1:**
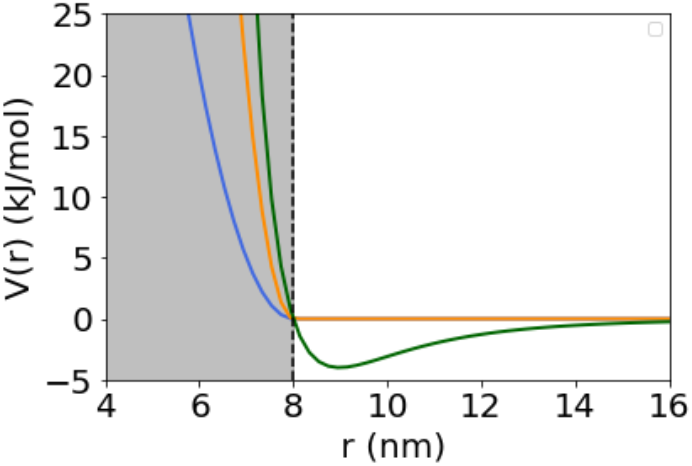
Interaction potentials used in simulations. The half-harmonic potentials with *κ*=10 kJ/mol/nm^2^ (*blue* line) and *κ*=40 kJ/mol/nm^2^ (*orange* line) and the LJ potential with *ϵ*=4 kJ/mol (*green* line) are shown. The vertical dotted line (in *black*) at *r*=8 nm indicates the contact distance for species *A* and *D*.

The system evolution obeys a reaction-diffusion formalism in which the particles make a Brownian diffusion step followed by a reaction step. The Brownian diffusion follows the overdamped Markovian Langevin dynamics:[52]

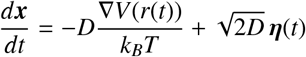

where ***x***(*t*) denotes the position of a particle with diffusion constant *D* (in water) under the influence of potential *V*(*r*) and **η**(*t*) is delta-correlated and Gaussian distributed noise in time such that

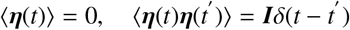

In simulations species *A, B* and *C* do not self-interact, but interact with the crowders via pair-wise forces implying that the concentration of these species can be obtained by their number densities. In order to avoid clustering of crowders, a self-repulsive harmonic potential with force constant *k*=10 kJ/mol/nm^2^ was added in all the simulations having a single or a mixed species of crowders. The system parameters chosen for simulations are listed in Table 1.

All the simulations are modeled as diffusion-limited reactions (see Supporting Information for details). Each simulation was run for 90 *μ*s with a time step of Δ*t*=0.012 ns and particle numbers were saved every 100 steps. To ensure that the trajectories are equilibrated at 300 K, the first 20 *μ*s were removed. Error bars were estimated by running 6 to 8 independent trajectories at each volume fraction and calculating the root mean square deviation from the average.

## 3. Results

### Homogeneous species of crowders interacting via repulsive potential

Macromolecular crowding has been shown to promote protein assembly[3, 4, 24, 29, 34, 55, 56]. Since many commonly used macromolecular crowders (e.g. dextran, ficoll) are neutral inert species, a soft harmonic repulsive potential was used to model this type of crowder-protein interactions. To verify the stabilization by crowders, the equilibrium free energy change in association reaction ΔΔ*F*_*AB*_(*ϕ*) was obtained as a function of volume fraction of crowders for two crowder sizes (Fig. 2 *top* panel). As expected, stabilization increases with increasing volume fraction of crowders. Moreover, smaller crowders provide better stabilization as has been observed previously[24, 28, 55]. To compare the free energy difference obtained from simulation with the theory, the force constant for the harmonic repulsive potential was increased from 10 kJ/mol/nm^2^ to 40 kJ/mol/nm^2^. In the limit of infinite stiffness the crowders would behave as hard spheres, but for soft crowders used in the simulations an effective hard sphere radius was calculated (see Supporting Information). The results indicate that theoretical estimates employing both hard and soft crowders tend to overestimate the free energy difference (Fig. 2 *bottom* panel). It was also found that, at a certain volume fraction, free energy depends weakly on the value of the force constant (Fig. S1).

**Figure 2:**
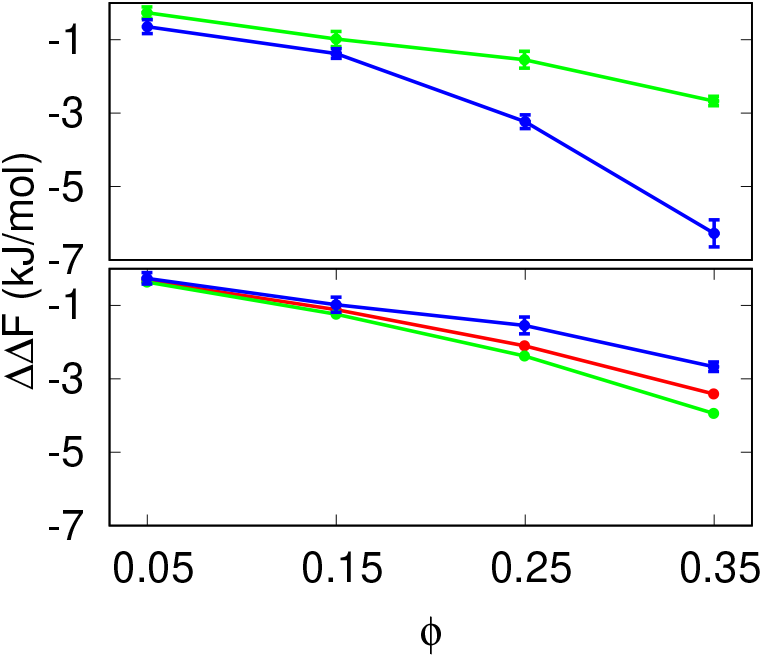
(*Top panel*) Free energy change ΔΔ*F* in presence of purely repulsive crowders showing that small crowders (*r*_*D*_=2.5 nm, *blue*) provide better stabilization than large crowders (*r*_*D*_=5 nm, *green*). (*Bottom panel*) Theoretical estimates of ΔΔ*F* employing both hard (*green*) and soft SPT (*red*) using *r*_*D*_=5 nm, showing overestimation of stabilization compared to *ReaDDy* simulation (*blue*).

The shape of the crowders control how much ‘free’ volume is available for the reaction to take place[57]. Consider a hard-sphere crowder dimer made up of two monomers of same radius. The dimer crowder provides more free volume for the reaction compared to the monomer crowder (see Supporting Information for details) and should disfavor the reaction (as the probability that the reactants come within the contact radius decreases). To test this in our simulations, crowder oligomers (dimer and tetramer) were modeled by joining monomeric particles of radius *r*_*d*_=2.5 nm with a harmonic bond of force constant *k*=100 kJ/mol/nm^2^.

The oligomers were kept linear by adding an harmonic angle potential with force constant *k*=80 kJ/mol/nm^2^. Fig. 3 shows small decrease in stabilization in going from monomer to dimer to tetramer indicating that shape change of the crowder does not make significant changes in the free energy. We note in passing that the above trend for hard-sphere oligomers cannot be readily applied for polymeric crowders at high concentration (in the semi-dilute regime) as the overlapping fraction of polymers is often significant under these circumstances[58].

**Figure 3:**
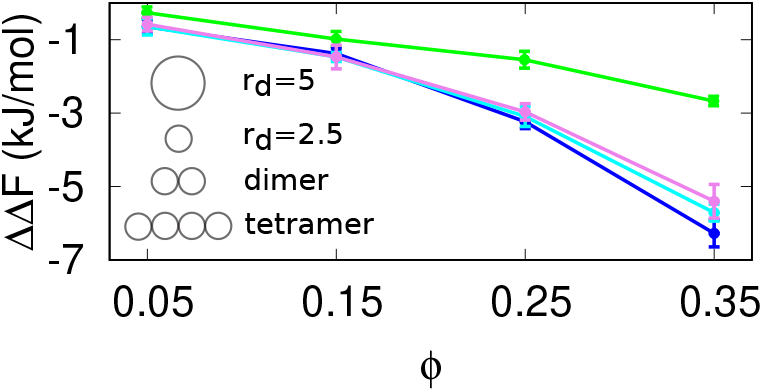
Free energy changes for crowders of varying shapes. Shown are crowder monomers with radii *r*_*D*_=5 nm (*green*), *r*_*D*_=2.5 nm (*blue*), dimer (*cyan*) and tetramer (*violet*). Crowder oligomers are formed by joining monomers with *r*_*D*_=2.5 nm by harmonic bond and angle potentials.

### Homogeneous species of crowders interacting via LJ potential

It has been postulated that weak attractive interactions by crowders can offset the excluded volume effect[24, 25, 37]. To incorporate attractive interactions we modeled crowder-protein interactions by a LJ potential with well-depth *ϵ*=4 kJ/mol (1.6 *k*_*B*_*T*) such that the repulsive part of the LJ potential matches closely with a purely repulsive potential with force constant k=40 kJ/mol/nm^2^. It is to be noted that *ϵ*=1.6 *k*_*B*_*T* gives a moderately strong LJ potential and is in the range of values used for same parameter in other works [30, 44]. Fig. 4 compares the free energy gain in the reaction from the two potentials. For LJ potential the reaction becomes less favorable with increasing volume fraction of crowders, showing a trend opposite to that observed for purely repulsive crowders. This behavior can be intuitively understood as follows: The LJ potential is long-ranged compared to the purely repulsive potential. At lowest volume fraction, the long range attraction between species *A, B* and crowder *D* increases the probability to make contacts between *A* and *B* and hence favors the forward reaction so that the reaction free energy is negative. The short range repulsive potential has negligible effect at lowest volume fraction. However, when *A* and *B* come in close contact and react, two particles are replaced by one *C* particle. The enthalpic cost of this replacement is negative for purely repulsive interactions, but positive for LJ interactions. Hence, with increasing volume fraction, free energy gain in the forward reaction increases for repulsive interactions, but decreases for LJ interactions. A weak LJ potential with *ϵ*=1 kJ/mol (0.4 *k*_*B*_*T*) showed stabilization with increasing volume fraction of the crowder species similar to the purely repulsive potential (data not shown). Following this trend, it seems likely that a stronger attraction (modeled by the well-depth of the LJ potential) by crowders can indeed disfavor the product formation and counter-balance the stabilization due to the repulsive part in a mixture of crowders.

**Figure 4:**
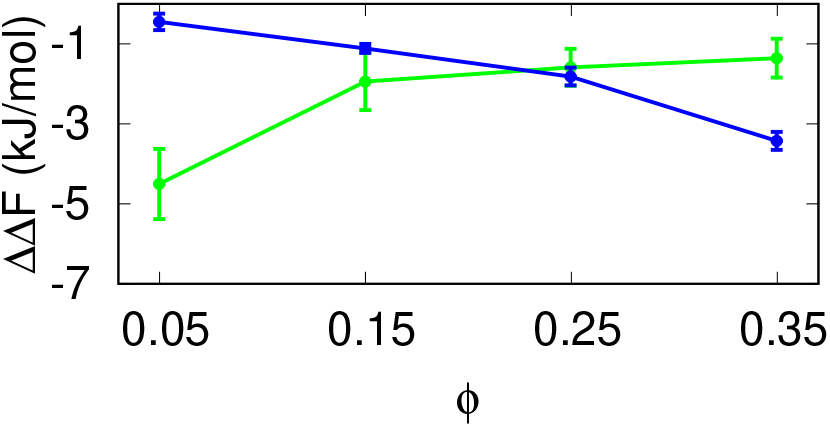
Change of free energy ΔΔ*F* for purely repulsive potential (*blue*) with k=40 kJ/mol/nm^2^ and attractive LJ potential (*green*) with *ϵ*=4 kJ/mol (1.6 *k*_*B*_*T*) for *r*_*D*_=5 nm.

### Mixed species of crowders

It remains unclear whether free energy change in presence of two or more crowders is additive of the free energy changes produced by the individual crowders. Several studies including computer simulations have reported that the effect of mixed macomolecular crowding is non-additive[20, 50, 51, 55], whereas other simulations showed that the free energy gain follows an additive ‘ansatz’[59, 60]. To verify the influence of mixed crowding conditions we prepared two different types of crowder mixtures: 1) Mixtures of crowders of two different sizes interacting via same (repulsive) potential and 2) Mixtures of two crowders of same sizes interacting via two different potentials.

#### Different sizes

Consider the reaction *A*+ *B* ⇌ *C* taking place at fixed volume fraction of crowders. Let ΔΔ*F_d_*_1_ and ΔΔ*F_d_*_2_ are the free energy changes in presence of crowders of radius d1 and d2, respectively. We prepared two crowder mixtures with relative concentration *d*1(0.75) : *d*2(0.25) and *d*1(0.25) : *d*2(0.75) and tested the additivity criteria as

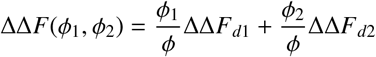

where, *ϕ*_1_ and *ϕ*_2_ are the relative concentrations of crowders in a mixture such that *ϕ*_1_ + *ϕ*_2_ = *ϕ*. Fig. 5 compares the free energy gain from simulations in a mixed crowder solution with that obtained using the additivity criteria. The results indicate that, given the simulation conditions, the additivity criteria remains valid for the entire range of volume fraction of crowders.

**Figure 5:**
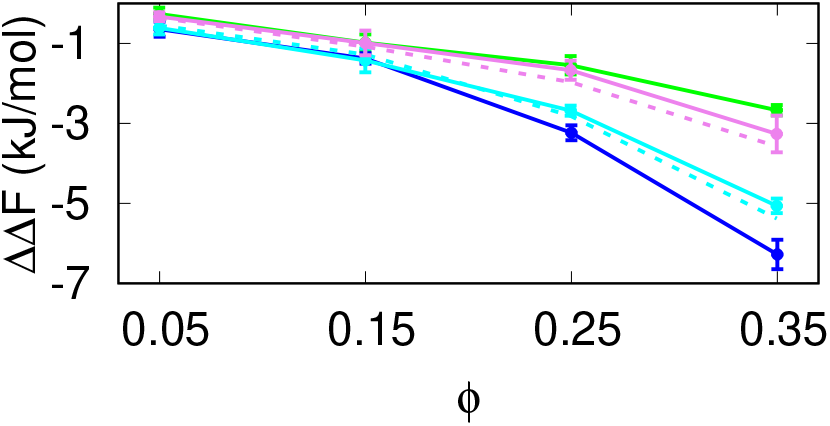
Stabilization in presence of mixed crowders of different sizes. Shown are the free energy changes for single type of crowders with *d*1=5 nm (*green*) and *d*2=2.5 nm (*blue*) and mixtures of them with relative concentration *d*1(0.25) : *d*2(0.75) (*cyan*) and *d*1(0.75) : *d*2(0.25) (*violet*). The dotted lines indicate the free energy changes calculated using the additivity ansatz.

#### Different interaction potentials

Next we consider two spherical crowders of same size, but each type of crowder species interacts with reactants and product with different potentials. Let the two types of crowders denoted by *R* and *L* interact via harmonic repulsive and LJ forces, respectively, with particles *A, B, C*. By preparing two crowder mixtures with relative concentration *R*(0.25) : *L*(0.75) and *R*(0.75) : *L*(0.25) we again test the additivity ‘ansatz’ for free energy.

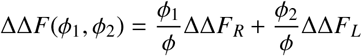

The results indicate that in a mixture where majority of particles interact by long-ranged LJ forces (Fig. 6, *top panel*), the free energy gain is clearly dominated by these particles and remain close to the free energy gain obtained for a solution of crowder particles interacting via LJ forces alone. In a binary mixture following these conditions, free energy is approximately additive at high volume fractions, but makes small deviation at low volume fraction. Interestingly, when majority of particles interact via purely repulsive forces (Fig. 6, *bottom panel*), free energy cost is far from additive. Instead, a minima in the free energy plot appears at volume fraction *ϕ* = 0.15, which is not seen for solution of crowders interacting by purely repulsive forces alone. To understand it further, we plotted the radial distribution function *g*(*r*) of *A* particles around the crowder species *L* for various *ϕ*, where *g*(*r*) was normalized with respect to the number density of *A* particles. It was found that the first peak of *g*(*r*) goes higher with increasing *ϕ*, implying that the probability of finding *A* particles around crowder *L* increases. Thus, the reaction becomes dominated by LJ forces at higher *ϕ*, while at low *ϕ* the repulsive species *R*, present in large volume fraction, plays the dominant role.

**Figure 6:**
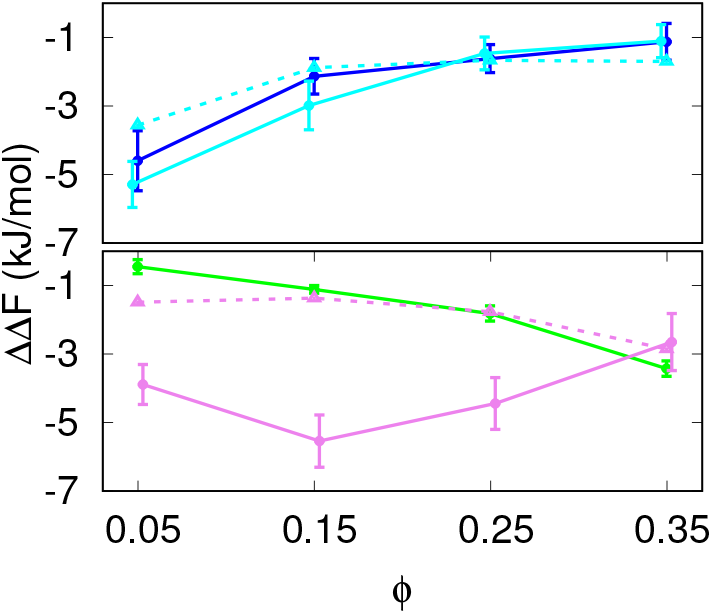
Stabilization in presence of mixed crowders interacting via different potentials. *(Top panel)* Free energy changes for single species of attractive *L* crowders (*blue*) and a binary mixture with relative concentration *R*(0.25) : *L*(0.75) (*cyan*). *(Bottom panel)* Free energy changes for single species of repulsive *R* crowders (*green*) and a binary mixture with relative concentration *R*(0.75) : *L*(0.25) (*violet*). The dotted lines indicate the free energy changes calculated using the additivity ansatz.

## 4. Discussion

*In-cell* association reactions occur in many different microenvironments with a variety of factors like crowding, confinement and adsorption regulating the thermo-dynamic reactivity. Since neither *in vivo* experiments nor computational models of cell-like environments can faithfully capture the complexity and heterogeneity of physiological media[35], we proposed a ‘bottom-up’ approach in this work, where complexity and heterogeneity of the system is increased in a step-wise manner to understand the crowding effects from simple to mixed environments. Although the results are qualitative in nature, it is important to understand the general principles of crowding influences from different sources of heterogeneity in a medium systematically, which is reported only by few studies [24, 30].

**Figure 7:**
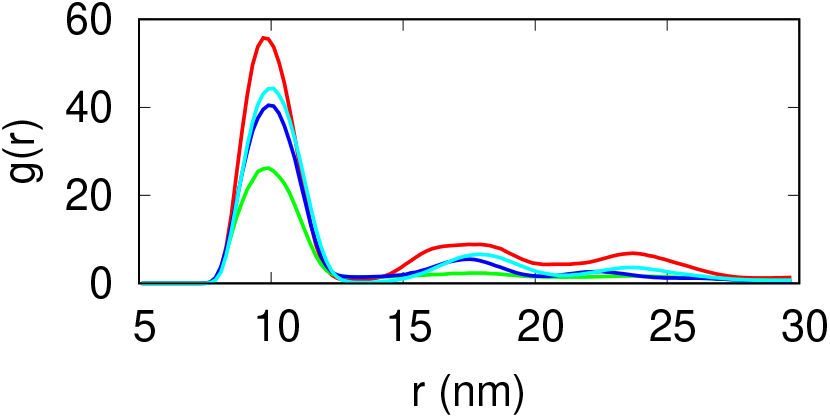
Radial distribution function of *A* particles around the crowder species *L* in a binary mixture with relative concentration *R*(0.75) : *L*(0.25). The total volume fraction is: *ϕ* = 0.05 (*green*), *ϕ* = 0.15 (*blue*) and *ϕ* = 0.25 (*cyan*). The *red* trace indicates the radial distribution function for a single species of LJ crowders at *ϕ* = 0.05.

In many simulation studies [29, 55] the transfer free energy 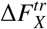 is estimated by calculating the excess chemical potential to put *one* particle of species *X* from infinitely dilute to crowded medium. This approach implicitly assumes that the crowded medium does not consist of any *X* particle and neglects self-interaction. The simulations presented here also make the same assumption by neglecting self-interaction for particle species *A, B* and *C*. However, for finite-sized particles self-interaction can be significant which introduces a correction factor to the association constant [61]. We plan to report on this in a separate study.

It was found that in a binary mixture of crowders free energy is additive of the same produced by individual components if protein interaction with both crowder species is of the same type (soft-repulsive) but not if the nature of interaction is different. It is to be noted here that in both cases the self- and hetero-interactions between crowders are taken to be repulsive. To verify if mutual attraction between crowders species can affect additivity of free energy, simulations were performed with a LJ potential (well-depth *ϵ*=4 kJ/mol) acting between species *D* and *E*. It was found that the free energy still remains approximately additive (see Fig.S2) in contrast with predictions made in earlier works [32, 59].

The main findings of this study are the following: 1) Crowding can both stabilize or destabilize a reaction depending on the nature of interaction with the reactants and products, 2) The shape of the crowder has marginal influence on reaction thermodynamics, and 3) The effect of mixed crowding environment depends on how we prepare the mixed solution. It is to be noted that, unlike the shape of the crowder, shape of the product in binary association can have a large influence in crowding-induced stabilization[33]. The advantage of the *ReaDDy* framework is that complex polymer and protein topologies can be modeled with a faithful description of inter-particle interactions. However, quantitative comparison with experiments (such as Ref.[50]) is a computationally demanding task and will be considered in a future study.

## Supporting information

Calculation of transfer free energy using SPT, equivalent hard sphere radius of soft particles, dependence of free energy on the force constant for the purely repulsive potential (Figure S1), calculation of free volume for crowders of various shapes, additivity of free energy for attractive hetero-interaction between crowders (Figure S2), simulation of well-mixed diffusion limited reaction in *ReaDDy.*

## References

[1] G. Schreiber, G. Haran, H.-X. Zhou, Fundamental aspects of protein-protein association kinetics, Chem. Rev. 109 (3) (2009) 839–860.

[2] M. Sakono, T. Zako, Amyloid oligomers: formation and toxicity of a*β* oligomers, FEBS J. 277 (6) (2010) 1348–1358.

[3] H.-X. Zhou, G. Rivas, A. P. Minton, Macromolecular crowding and confinement: biochemical, biophysical, and potential physiological consequences, Annu. Rev. Biophys. 37 (2008) 375–97.

[4] I. Kuznetsova, K. Turoverov, V. Uversky, What macro-molecular crowding can do to a protein, Int. J. Mol. Sci. 15 (12) (2014) 23090–23140.

[5] S. Shahid, M. I. Hassan, A. Islam, F. Ahmad, Size-dependent studies of macromolecular crowding on the thermodynamic stability, structure and functional activity of proteins: in vitro and in silico approaches, Biochim. Biophys. Acta 1861 (2) (2017) 178–197.

[6] G. Rivas, A. P. Minton, Toward an understanding of biochemical equilibria within living cells, Biophys. Rev. 10 (2) (2018) 241–253.

[7] M. Sarkar, C. Li, G. J. Pielak, Soft interactions and crowding, Biophys. Rev. 5 (2) (2013) 187–194.

[8] A. Politou, P. A. Temussi, Revisiting a dogma: the effect of volume exclusion in molecular crowding, Curr. Opin. Struct. Biol. 30 (2015) 1–6.

[9] D. Gnutt, S. Ebbinghaus, The macromolecular crowding effect-from in vitro into the cell, Biol. Chem. 397 (1) (2016) 37–44.

[10] D. Guin, M. Gruebele, Weak chemical interactions that drive protein evolution: crowding, sticking, and quinary structure in folding and function, Chem. Rev. 119 (18) (2019) 10691–10717.

[11] A. G. Ogston, The spaces in a uniform random suspension of fibres, Trans. Faraday Soc. 54 (1958) 1754–1757.

[12] T. C. Laurent, The interaction between polysaccharides and other macromolecules. 5. the solubility of proteins in the presence of dextran, Biochem. J. 89 (1963) 253–7.

[13] A. P. Minton, Excluded volume as a determinant of macromolecular structure and reactivity, Biopolymers 20 (10) (1981) 2093–2120.

[14] O. G. Berg, The influence of macromolecular crowding on thermodynamic activity: Solubility and dimerization constants for spherical and dumbbell-shaped molecules in a hard-sphere mixture, Biopolymers 30 (11-12) (1990) 1027–1037.

[15] S. B. Zimmerman, S. O. Trach, Estimation of macro-molecule concentrations and excluded volume effects for the cytoplasm of Escherichia coli, J. Mol. Biol. 222 (3) (1991) 599–620.

[16] R. R. Gabdoulline, R. C. Wade, Simulation of the diffusional association of barnase and barstar, Biophys. J. 72 (5) (1997) 1917–29.

[17] Y. Qu, C. L. Bolen, D. W. Bolen, Osmolyte-driven contraction of a random coil protein, Proc. Natl. Acad. Sci. USA 95 (16) (1998) 9268–73.

[18] H.-X. Zhou, Protein folding and binding in confined spaces and in crowded solutions, J. Mol. Recognit. 17 (5) (2004) 368–75.

[19] M. S. Cheung, D. Klimov, D. Thirumalai, Molecular crowding enhances native state stability and refolding rates of globular proteins, Proc. Natl. Acad. Sci. USA 102 (13) (2005) 4753–8.

[20] J. Batra, K. Xu, H.-X. Zhou, Nonadditive effects of mixed crowding on protein stability, Proteins 77 (1) (2009) 133–8.

[21] A. C. Miklos, M. Sarkar, Y. Wang, G. J. Pielak, Protein crowding tunes protein stability, J. Am. Chem. Soc. 133 (18) (2011) 7116–7120.

[22] R. Harada, Y. Sugita, M. Feig, Protein crowding affects hydration structure and dynamics, J. Am. Chem. Soc. 134 (10) (2012) 4842–4849.

[23] Y. Wang, M. Sarkar, A. E. Smith, A. S. Krois, G. J. Pielak, Macromolecular crowding and protein stability, J. Am. Chem. Soc. 134 (40) (2012) 16614–16618.

[24] Y. C. Kim, J. Mittal, Crowding induced entropy-enthalpy compensation in protein association equilibria, Phys. Rev. Lett. 110 (20) (2013) 208102.

[25] M. Senske, L. Törk, B. Born, M. Havenith, C. Herrmann, S. Ebbinghaus, Protein stabilization by macromolecular crowding through enthalpy rather than entropy, J. Am. Chem. Soc. 136 (25) (2014) 9036–9041.

[26] N. F. Dupuis, E. D. Holmstrom, D. J. Nesbitt, Molecular-crowding effects on single-molecule rna folding/unfolding thermodynamics and kinetics, Proc. Natl. Acad. Sci. USA 111 (23) (2014) 8464–9.

[27] S. K. Mukherjee, S. Gautam, S. Biswas, J. Kundu, P. K. Chowdhury, Do macromolecular crowding agents exert only an excluded volume effect? a protein solvation study, J. Phys. Chem. B 119 (44) (2015) 14145–14156.

[28] K. A. Sharp, Analysis of the size dependence of macro-molecular crowding shows that smaller is better, Proc. Natl. Acad. Sci. USA 112 (26) (2015) 7990–7995.

[29] T. Ando, I. Yu, M. Feig, Y. Sugita, Thermodynamics of macromolecular association in heterogeneous crowding environments: Theoretical and simulation studies with a simplified model, J. Phys. Chem. B 120 (46) (2016) 11856–11865.

[30] T. Hoppe, A. P. Minton, Incorporation of hard and soft protein–protein interactions into models for crowding effects in binary and ternary protein mixtures. comparison of approximate analytical solutions with numerical simulation, J. Phys. Chem. B 120 (46) (2016) 11866–11872.

[31] M. Feig, I. Yu, P.-H. Wang, G. Nawrocki, Y. Sugita, Crowding in cellular environments at an atomistic level from computer simulations, J. Phys. Chem. B 121 (34) (2017) 8009–8025.

[32] A. P. Minton, Explicit incorporation of hard and soft protein–protein interactions into models for crowding effects in protein mixtures. 2. effects of varying hard and soft interactions upon prototypical chemical equilibria, J. Phys. Chem. B 121 (22) (2017) 5515–5522.

[33] A. J. Guseman, G. M. P. Goncalves, S. L. Speer, G. B. Young, G. J. Pielak, Protein shape modulates crowding effects, Proc. Natl. Acad. Sci. USA 115 (2018) 10965–10970. doi:10.1073/pnas.1810054115.

[34] S. B. Zimmerman, A. P. Minton, Macromolecular crowding: Biochemical, biophysical, and physiological consequences, Annu. Rev. Biophys. Biomol. Struct. 22 (1993) 27–65.

[35] G. Rivas, A. P. Minton, Macromolecular crowding in vitro, in vivo, and in between, Trends Biochem. Sci. 41 (11) (2016) 970–981.

[36] R. Harada, N. Tochio, T. Kigawa, Y. Sugita, M. Feig, Reduced native state stability in crowded cellular environment due to protein-protein interactions, J. Am. Chem. Soc. 135 (9) (2013) 3696–3701.

[37] M. Sarkar, J. Lu, G. J. Pielak, Protein crowder charge and protein stability, Biochemistry 53 (10) (2014) 1601–1606.

[38] W. G. McMillan Jr, J. E. Mayer, The statistical thermodynamics of multicomponent systems, J. Chem. Phys. 13 (7) (1945) 276–305.

[39] I. L. Shulgin, E. Ruckenstein, Various contributions to the osmotic second virial coefficient in protein-water-cosolvent solutions, J. Phys. Chem. B 112 (46) (2008) 14665–14671.

[40] G. T. Weatherly, G. J. Pielak, Second virial coefficients as a measure of protein-osmolyte interactions, Protein Sci. 10 (1) (2001) 12–16.

[41] C. Li, G. J. Pielak, Using nmr to distinguish viscosity effects from nonspecific protein binding under crowded conditions, J. Am. Chem. Soc. 131 (2009) 1368–1369. doi:10.1021/ja808428d.

[42] Y. Wang, C. Li, G. J. Pielak, Effects of proteins on protein diffusion, J. Am. Chem. Soc. 132 (27) (2010) 9392–7.

[43] D. Hall, A. P. Minton, Macromolecular crowding: qualitative and semiquantitative successes, quantitative challenges, Biochimica et Biophysica Acta 1649 (2) (2003) 127–139.

[44] L. Sapir, D. Harries, Origin of enthalpic depletion forces, J. Phys. Chem. Lett. 106 (2014) 671a. doi:10.1016/j.bpj.2013.11.3719.

[45] H. Reiss, H. L. Frisch, J. L. Lebowitz, Statistical mechanics of rigid spheres, J. Chem. Phys. 31 (2) (1959) 369–380.

[46] J. L. Lebowitz, E. Helfand, E. Praestgaard, Scaled particle theory of fluid mixtures, J. Chem. Phys. 43 (3) (1965) 774.

[47] M. A. Cotter, Hard spherocylinders in an anisotropic mean field: A simple model for a nematic liquid crystal, J. Chem. Phys. 66 (3) (1977) 1098–1106.

[48] H.-X. Zhou, S. Qin, Simulation and modeling of crowding effects on the thermodynamic and kinetic properties of proteins with atomic details, Biophys. Rev. 5 (2) (2013) 207–215.

[49] H.-X. Zhou, Effect of mixed macromolecular crowding agents on protein folding, Proteins 72 (4) (2008) 1109–13.

[50] S. Biswas, J. Kundu, S. K. Mukherjee, P. K. Chowdhury, Mixed macromolecular crowding: A protein and solvent perspective, ACS Omega 3 (4) (2018) 4316–4330.

[51] S. Shahid, F. Ahmad, M. I. Hassan, A. Islam, Mixture of macromolecular crowding agents has a non-additive effect on the stability of proteins, Appl. Biochem. Biotechnol. 188 (4) (2019) 927–941.

[52] J. Schöneberg, F. Noé, Readdy-a software for particle-based reaction-diffusion dynamics in crowded cellular environments, PLoS One 8 (9) (2013) e74261.

[53] M. B. Dolinska, P. T. Wingfield, Y. V. Sergeev, *β*b1-crystallin: thermodynamic profiles of molecular interactions, PloS one 7 (1) (2012) e29227.

[54] J. S. Kim, A. Yethiraj, Crowding effects on protein association: effect of interactions between crowding agents, J. Phys. Chem. B 115 (2) (2011) 347–353.

[55] S. Qin, H.-X. Zhou, Atomistic modeling of macromolecular crowding predicts modest increases in protein folding and binding stability, Biophys. J. 97 (1) (2009) 12–9.

[56] M. Jiao, H.-T. Li, J. Chen, A. P. Minton, Y. Liang, Attractive protein-polymer interactions markedly alter the effect of macromolecular crowding on protein association equilibria, Biophys. J. 99 (3) (2010) 914–923.

[57] A. Samiotakis, M. S. Cheung, Folding dynamics of trp-cage in the presence of chemical interference and macro-molecular crowding. i, J. Chem. Phys. 135 (17) (2011) 175101.

[58] N. Kozer, Y. Y. Kuttner, G. Haran, G. Schreiber, Protein-protein association in polymer solutions: from dilute to semidilute to concentrated, Biophys. J. 92 (6) (2007) 2139–2149.

[59] Y. C. Kim, R. B. Best, J. Mittal, Macromolecular crowding effects on protein–protein binding affinity and specificity, J. Chem. Phys. 133 (20) (2010) 11B608.

[60] A. Kudlay, M. S. Cheung, D. Thirumalai, Influence of the shape of crowding particles on the structural transitions in a polymer, J. Phys. Chem. B 116 (2012) 8513–8522. doi:10.1021/jp212535n.

[61] D. H. De Jong, L. V. Schäfer, A. H. De Vries, S. J. Marrink, H. J. Berendsen, H. Grubmüller, Determining equilibrium constants for dimerization reactions from molecular dynamics simulations, J. Comput. Chem. 32 (9) (2011) 1919–1928.

