## Supporting Information for "Modeling Protein Association from Homogeneous to Mixed Environments: A Reaction-Diffusion Dynamics Approach"

### 1 Calculation of transfer free energy using SPT.

In SPT, crowders are modeled as hard convex particles and the free energy to transfer any particle $X$ ($\Delta F_X^{tr}$) from a dilute solution to a crowded environment is calculated. Consider a homogeneous solution made of spherical molecules of type $D$ (crowders) having radius $r_D$. The free energy ($\Delta F_A^{tr}$) of transferring a spherocylindrical molecule $A$ having cylindrical radius $r_A$ and cylindrical length $l_A$ in this solution can be expressed as[?]

$$\Delta F_A^{tr} = RT\left[-ln(1-\phi) + a_1 Q + a_2 Q^2 + a_3 Q^3\right],$$

where $\phi$ denotes the volume fraction of the fluid spheres, and

$$\begin{aligned}
Q &= \frac{\phi}{1-\phi},\\
a_1 &= r^3 + 3r^2 + 3r + 1.5l(r^2 + 2r + 1),\\
a_2 &= 1.5(2r^3 + 3r^2) + 4.5l(r^2 + r),\\
a_3 &= 3r^3 + 4.5lr^3.
\end{aligned}$$

$r$ and $l$ are scaled parameters defined by

$$\begin{aligned}
r &\equiv \frac{r_A}{r_D},\\
l &\equiv \frac{l_A}{2r_A},
\end{aligned}$$

### 2 Equivalent hard sphere radius of soft particles.

For a solution of particles interacting via soft potentials, the volume excluded $V_{ex}$ can be obtained by negative of the volume integral of the Mayer f-function[?] as

$$V_{ex} = -\int\left(\exp\left[-\frac{U(r)}{kT}\right] - 1\right) d^3r$$

where $U(r)$ denotes the interaction potential between a pair of particles. Assuming the interacting particles are of the same type, the equivalent hard sphere radius $r_{hard}$ is defined using

$$2r_{hard} = \left(\frac{3}{4\pi}V_{ex}\right)^{\frac{1}{3}}$$

### 3 Dependence of free energy on the force constant for the purely repulsive potential.

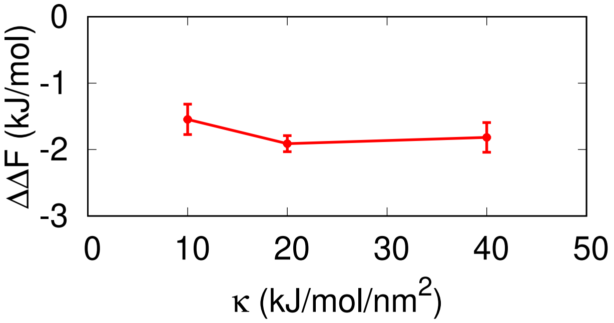


Figure S1: Free energy change $\Delta\Delta F$ for purely repulsive crowders at a constant volume fraction $\phi = 0.25$ for various force constant $\kappa$.

### 4 Calculation of free volume for crowders of various shapes.

If $\phi$ denotes the volume fraction of crowders in a box of volume V, then the volume accessible (free) for the reaction is

$$V_f = V\,(1-\phi)$$

In the present model species $A, B, C$ do not interact with each other or themselves, but can interact with $D$. Hence the free volume reduces to

$$V_f = \begin{cases} V - n_D \times \frac{4}{3}\pi\left(r_D + r_{A/B}\right)^3 & \text{for A/B} \\ V - n_D \times \frac{4}{3}\pi\left(r_D + r_C\right)^3 & \text{for C} \end{cases}$$

where $n_D$ is the number of crowder species $D$ and $r_X$ denotes the radius of species $X$ ($X \in (A, B, C, D)$). For crowder dimers interacting with finite sized particles (A, B, C), a part of the crowder surface area becomes inaccessible due to dimerization (Fig. S2) and the volume excluded by each crowder particle (for species $A$) is given by

$$\int_0^{2\pi}\int_0^{\pi-\theta_0}\int_0^{R} r^2 \sin\theta dr d\theta d\phi = \frac{2\pi R^3}{3}\left(1+\cos\theta_0\right)$$

where $R = r_D + r_A$.

Hence, the free volume for crowder dimers become

$$V_f = V - n_D \times \frac{2\pi R^3}{3}\left(1+\frac{r_D}{R}\right)$$

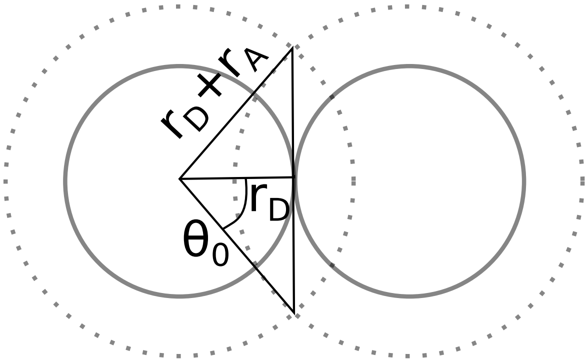


Figure S2: Schematic diagram for calculation of volume excluded by a dimer of species $D$ while interacting with species $A$ via hard-sphere potential.

Since, $\frac{r_D}{R} < 1$, we have $V_f^{dimer} > V_f^{monomer}$.

### 5 Additivity of free energy for attractive hetero-interaction between crowders.

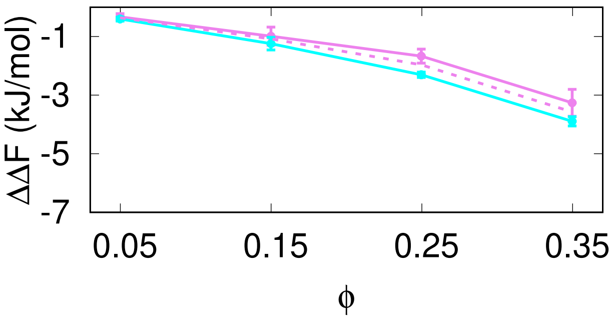


Figure S3: Free energy change for mixture of crowder species with $d1 = 5$ nm and $d2 = 2.5$ nm with relative concentration $d1(0.75) : d2(0.25)$. Shown are the traces for purely repulsive (*violet*) and LJ (*cyan*) hetero-interactions between the crowders. The dotted line indicates the free energy change calculated using the additivity ansatz on single type of crowder species.

### 6 Simulation of well-mixed diffusion limited reaction.

In *ReaDDy*, the association reaction is modeled as

$$A + B \underset{k_{off}}{\overset{k_{enc}}{\rightleftarrows}} [AB] \xrightarrow{\lambda_{on}} C$$

where A and B come in contact with each other to form an encounter complex [AB] with a rate $k_{enc}$ and reacts with a rate $\lambda_{on}$ to form a product C. The dissociation rate constant is given by $k_{off}$. To simulate a diffusion-limited reaction we must have $\lambda_{on} \gg k_{off}$. Secondly, the condition of well-mixing demands that diffusion must be faster than the reaction time scale, that is, $\frac{r^2}{D} \ll \lambda_{on}^{-1}$. Finally, the time step $\Delta t$ for integration the equation of motion should also be smaller than $\frac{r^2}{D}$. To satisfy these conditions in our simulations, we have chosen $\lambda_{on} = 10^{-3}$ ns$^{-1}$, $k_{off} = 10^{-5}$ ns$^{-1}$ and $\Delta t = 10^{-4} \times \frac{r_A^2}{D_A} \approx 0.012$ ns.
